# Efficient murine cardiac phenotyping by combining synchrotron-based phase-contrast micro-CT, histology, immunofluorescence and spatial transcriptomics

**DOI:** 10.64898/2026.04.24.720617

**Authors:** Niccolò Peruzzi, Ayse Ceren Mutgan, Kinga I. Gawlik, Timothy J. Mead, Tuva Jerlhagen Forsgren, Elin Lagervall, Till Dreier, Robin Krüger, Elna Lampei, Jenny Romell, Rapolas Spalinskas, Madeleine Durbeej-Hjalt, Suneel S. Apte, Martin Bech, Anne Bonnin, Katarina Tiklova, Karin Tran-Lundmark

## Abstract

Congenital heart disease is commonly studied using genetically modified mouse models, but characterization of complex three-dimensional (3D) cardiac anomalies remains technically challenging. Histology of fixed paraffin-embedded samples, typically used for structural and molecular analysis, is limited to two dimensions (2D). Synchrotron radiation-based phase-contrast micro-computed tomography (SRPC-µCT) enables rapid, high-resolution, 3D imaging but had yet to be fully integrated with molecular tissue analysis. Importantly, SRPC-µCT is nondestructive and compatible with paraffin-embedded tissue, potentially allowing integration with downstream molecular analyses such as multiplexed spatial transcriptomic profiling on tissue sections.

Here, we present a pipeline combining SRPC-µCT with complementary molecular approaches for rapid and accurate phenotyping of mouse hearts and for investigating disease mechanisms. We demonstrate the successful integration of high-resolution 3D imaging with spatial transcriptomics, fluorescent stainings, and histochemistry. Notably, all 2D modalities were applied sequentially to a single tissue section and registered within the 3D volume. This multimodal framework provides a powerful approach for linking structural and molecular information that is broadly applicable across biomedical research.

## 2. Introduction

Congenital heart disease (CHD) comprises diverse structural birth defects with a wide spectrum of functional impact, often deadly, affecting almost 2% of human live births and ranking among the top 8 leading causes of death in children younger than 1 year ^1^. As such, the etiology, mechanisms and management of CHD are subjects of widespread research. Genetically modified mouse models are routinely used in studies of CHD ^2^. Cardiac phenotyping in mice is, however, quite challenging because of the small size of a developing mouse heart: a typical embryonic day (E) 18.5 mouse heart is just ∼3 mm in diameter. The most commonly employed technique is histological assessment of formalin-fixed paraffin-embedded (FFPE) samples, which, however, provides only two-dimensional (2D) information and thus might miss important structural defects. Most methods to obtain three-dimensional (3D) information with sufficient resolution are either time-consuming (e.g. histological serial sectioning followed by digital 3D reconstruction), require specific sample preparation (e.g. light sheet microscopy of cleared tissue), or cause complete destruction of the sample (e.g. high-resolution episcopic microscopy) ^3^.

In recent years, synchrotron radiation-based (SR) phase-contrast micro-computed tomography (PC-µCT) has emerged as a powerful tool for 3D assessment of tissue micro-structure *ex vivo*. This technique exploits the high coherence of synchrotron light to enhance image contrast in tomographic measurements of soft tissue without the need for staining agents. The high flux of synchrotron light also allows high-magnification microscopes in the detection system without having to substantially sacrifice scan speed or image quality. Lastly, a significant advantage of PC-µCT over other methods is that it does not require specific sample preparation and is nondestructive. The imaged tissue can therefore be sectioned and used for methods such as histology, immunohistochemistry and *in situ* hybridization for comparison and integration with the CT scan ^4–6^.

SRPC-µCT has successfully been used for imaging hearts from different animal models (mice ^7–12^, rats ^13–15^, rabbits ^14^), as well as for visualizing explanted human heart ^16–18^. While several of these studies have shown the potential of SRPC-µCT to provide morphological assessment of cardiac defects, the more recent focus has been on the cardiac micro-structure, exploiting the high resolution (< 1 µm) that can be achieved with SRPC-µCT. Because high resolution necessarily compromises the field-of-view (FoV) of a single scan, in most of these publications scans of large hearts have been achieved by acquiring multiple sub-volumes and digitally stitching them together. As a culmination of this approach, whole human adult hearts have recently been scanned with 20 µm voxel size, with a few selected areas imaged with as low voxel size as 2.26 µm ^19,20^. While the option to achieve µm-resolution in whole large organs is a significant advantage of SRPC-µCT compared to other microscopy methods, it also results in very long scan times (the two human adult heart scans in ^20^ are reported to have required ∼7 h each for the whole organ scans, plus ∼2 h for each of the higher resolution areas); combined with the low availability of synchrotron beam time, this use seems for now restricted to the creation of high quality atlases, or to studies with a limited number of samples. In the case of mouse fetal hearts, however, a single E18.5 heart can fit within the FoV of the ∼2 µm resolution configuration of most PC-µCT synchrotron beamlines; as such, a full scan could require only a few minutes without compromising on quality. The small size of mouse hearts would also be optimal when trying to perform PC-µCT of unstained samples in a laboratory setting, a method which has recently seen steady developments ^21–23^. Even though scan times with laboratory scanners are longer, easier accessibility compared to synchrotrons could lead to a wider implementation of PC-µCT into phenotyping routines, still retaining at the same time full compatibility with other methods such as traditional histology. While a significant part of this paper is dedicated to PC-µCT performed at synchrotrons, where large batches of samples could in principle be scanned in a very short time ^24^, some results from laboratory-based (LB) scanners, a custom made one and a commercial scanner capable of performing PC-µCT, will also be shown and discussed as a potential, more accessible, alternative.

Although some studies have hinted at how PC-µCT does not impair subsequent histological and molecular evaluation of the hearts, most existing papers have limited themselves to showing views of comparable areas, but have not precisely registered slices from the different techniques. Nevertheless, methods to achieve a 1-to-1 match have been already proved in literature for other types of tissue ^5,25–29^. A precise alignment of different modalities would provide structural context to the 2D observations enabled by histology and could furthermore serve as a basis for machine learning or deep learning approaches in the future. In this work, we present how PC-µCT can enable an accurate and fast phenotyping of fetal mouse hearts, and how an analysis pipeline could take form, with a particular focus on communication in multidisciplinary teams and on full integration of methods directed at understanding the molecular mechanisms behind different types of congenital heart disease. Methods for analysis of FFPE tissue have developed extremely fast over the last couple of years. It is now possible to study 5000 mRNAs at the same time in a single tissue section using *in situ* sequencing (ISS) with padlock probes and rolling-circle amplification (Xenium®). This allows targeted transcriptional profiling of RNA directly within tissue sections while preserving cellular spatial organization. To our knowledge, our study is the first to fully integrate spatial transcriptomics with SRPC-µCT. In addition, we explored the possibility to use the same tissue section for histology and immunofluorescence following ISS.

## 3. Results

In this section we present the results obtained at different stages of our analysis pipeline. The analysis was performed with a combination of different functions in the software programs FIJI ^30^ and Amira (Thermo Fisher Scientific, Waltham, Massachusetts, US). Parts of the pipeline are described in this section as necessary; more details are provided in the Methods section.

As a first step, we show how morphological information about the hearts can be extracted directly from the SRPC-µCT scans. Subsequently, we present an example in which we used this morphological information for cardiac phenotyping of an *Adamts6*-deficient mouse strain. We then introduce a pipeline for integration of 2D histological slides into 3D PC-µCT volumes and show how we used it to successfully combine SRPC-µCT with histochemical staining, fluorescent stainings, and spatial transcriptomics. The final section presents results from laboratory-based µCT systems and offers a qualitative comparison between them and the synchrotron results.

### 3.1. Morphology

A first assessment of the SRPC-µCT data consisted in scrolling through the volumes with orthogonal slices (Figure 1-A). A preliminary 3D visualization, without the need of any additional pre-processing, was also performed using a semi-transparent volume rendering (Figure 1-B); by adjusting transparency based on the image grayscale, the organs were visualized as distinct from the paraffin embedding in just a few seconds. This coarse visualization, together with the use of orthogonal slices, was employed to rapidly assess sample and image quality and to inform the subsequent physical sectioning of the samples (i.e. anticipating how the heart would be positioned while cutting through the block and deciding at roughly which depth slices should be extracted for staining).

**Figure 1.**
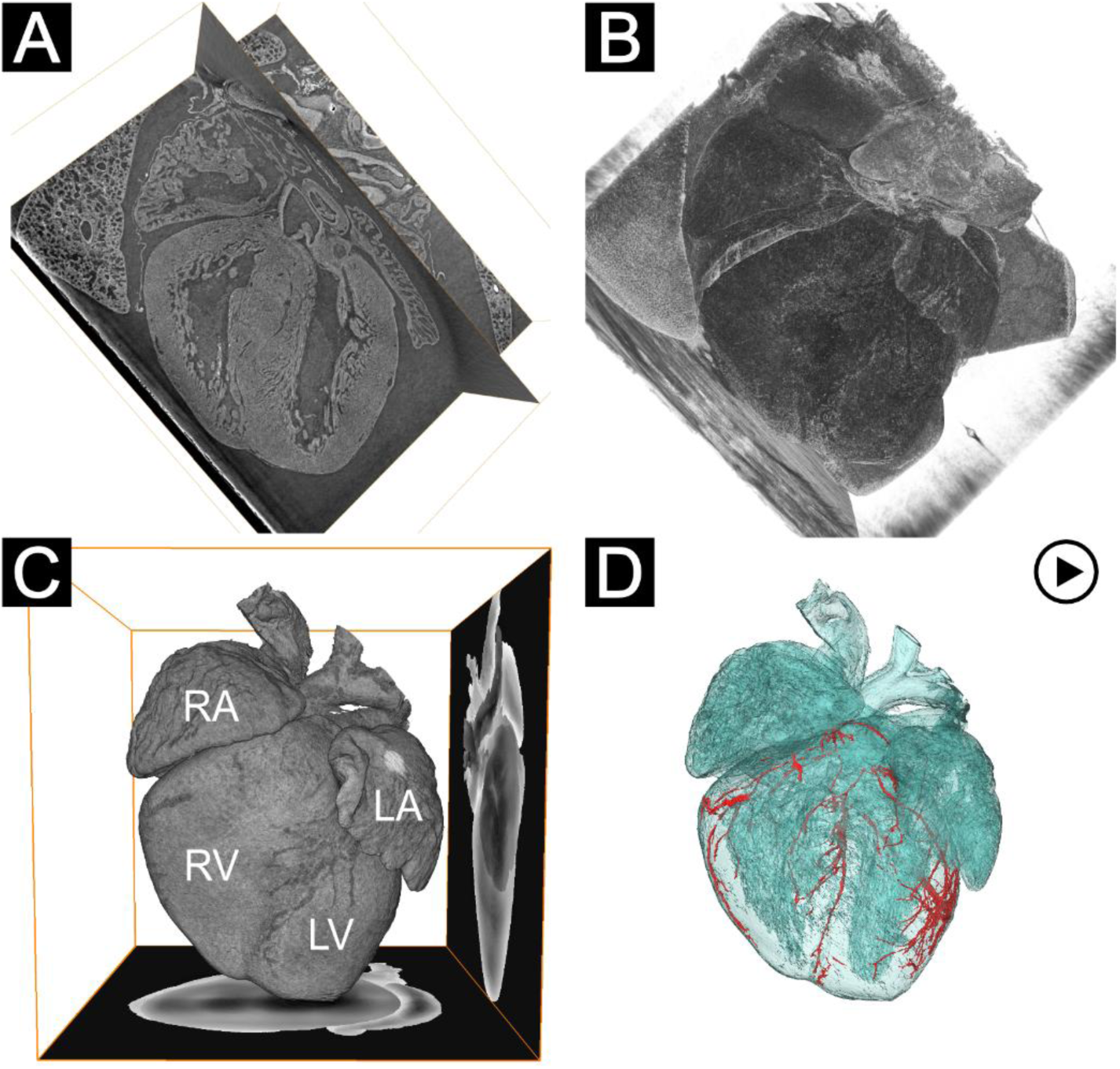
Overview of cardiac morphology assessment from SRPC-µCT data at increasing levels of complexity (and proportionately, time) in data segmentation. **A** – An immediate evaluation consists in digitally re-slicing the volume along arbitrary directions. Multiple slicing planes can be combined. **B** – A semi-transparent volume rendering can be directly applied to the data without any preprocessing. The overall structure of the heart is visible, but morphological details are hidden by the pericardium and other organs. **C** – After segmentation of the heart and removal of the pericardium, the heart structure is readily visible with a volume rendering. RA: right atrium, LA: left atrium, RV: right ventricle, LV: left ventricle. The rendering is in a cubic box with 4-mm-long sides, on two faces of which the segmented dataset is projected, to more easily visualize the heart size. **D** – The coronary circulation is segmented and shown as a red mesh surface rendering, together with a transparent mesh surface view of the heart to provide positional context. **Video-clip 1** illustrates the transition between the different visualization options.

As a next step of the analysis, the hearts were segmented from the rest of the tissue. The resulting cleaned segmentations allowed creation of 3D renderings of the hearts (Figure 1-C) that were used as a basis for all the other visualizations.

As a further example of what could be achieved from these datasets, a semi-automatic segmentation of the coronary circulation was performed. A mesh surface rendering of the coronary circulation is shown in red in Figure 1-D, together with a transparent mesh surface view of the heart to provide positional context.

A showcase of the different visualization options is provided in Video-clip 1. The 3D volume can be digitally sectioned along any arbitrary plane. While this can be performed immediately, without any segmentation, a volume rendering often aids in positioning the slices and provides a more immediate understanding of the 3D context. To facilitate evaluation by specialists with a medical background, standard echocardiography views can be digitally recreated. An example of this is provided in Figure 2, where the left parasternal long-axis views, commonly employed in echocardiography ^31^, were recreated on the PC-µCT data (Figure 2). Using the volume rendering shown in Figure 1-C as a reference (here colored in pink, to distinguish it from the grayscale slices), digital slices in the Amira 3D environment could be positioned to mimic a sweep from an echocardiography transducer. Slices matching the three different views are shown with a yellow border; the volume rendering can also be cut in correspondence of these slices to extract thin 3D slabs, as shown right below.

**Figure 2.**
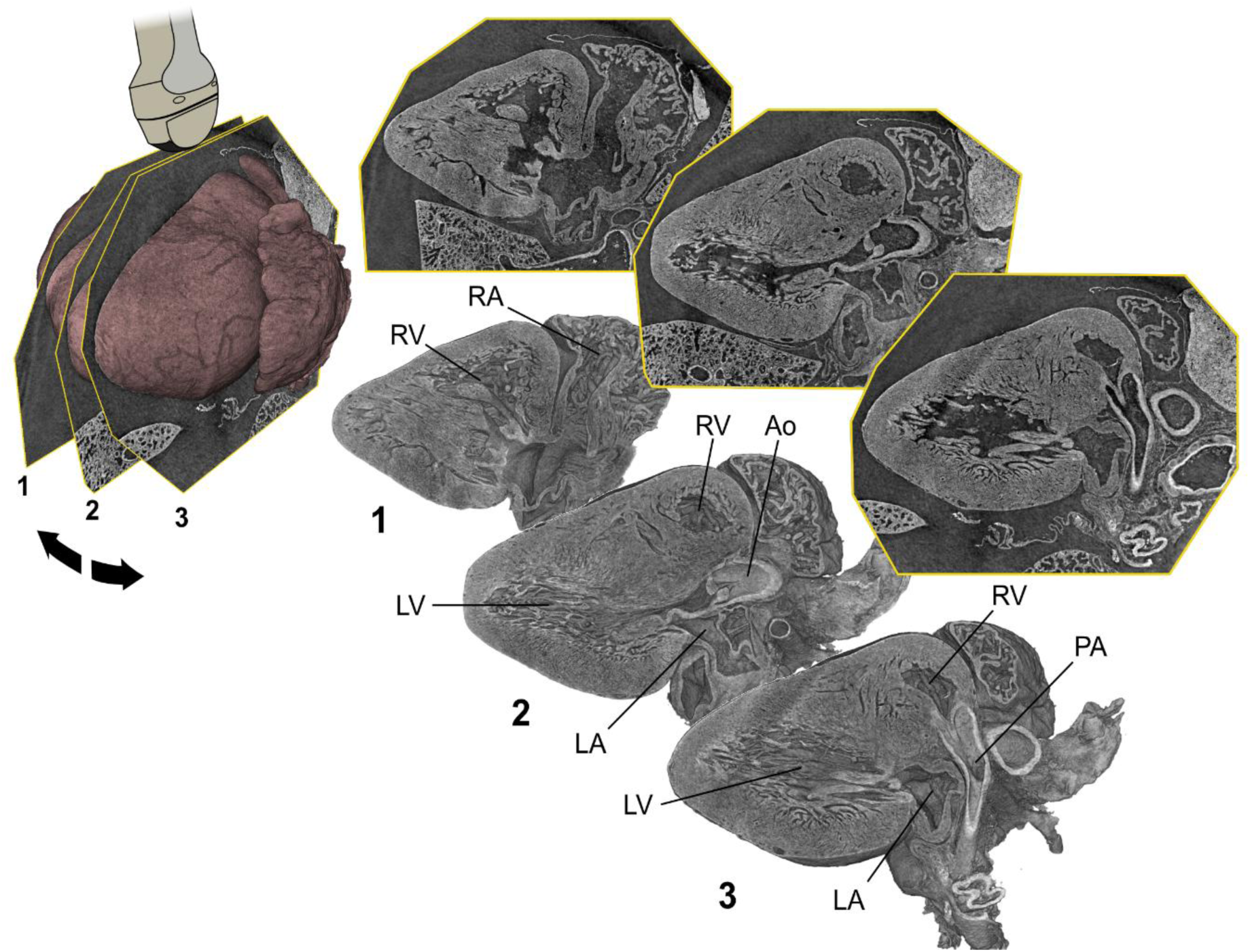
Standard echocardiography views can be digitally recreated to assist in evaluation. Here, left parasternal long-axis views (as described in Lai et al.) were created from the PC-µCT data. RA: right atrium, LA: left atrium, RV: right ventricle, LV: left ventricle, Ao: aorta, PA: pulmonary artery.

### 3.1. Potential for phenotyping

The structural information provided by PC-µCT can be used for fast and precise phenotyping. As an example, Figure 3 illustrates the case of an *Adamts6*-deficient mouse strain. *Adamts6*-deficient mice have previously been demonstrated to develop cardiac defects and other congenital anomalies ^32–34^. With PC-µCT, apical 5-chamber views of wild-type (Figure 3-A) and mutant (Figure 3-C) hearts were readily obtained and compared. In Figure 3, volume renderings similar to the ones shown in Figure 1-C and 2-B are used. For additional clarity, the luminal volumes of the cardiac chambers were also segmented from the 3D volumes and color-coded to distinguish laterality, with right-sided structures shown in blue and left-sided structures in red. In the case of the *Adamts6*-deficient mouse strain, a ventricular septal defect is recognizable (color-coded in purple for clarity). Figure 3-B shows a schematic of the defect being observed.

**Figure 3.**
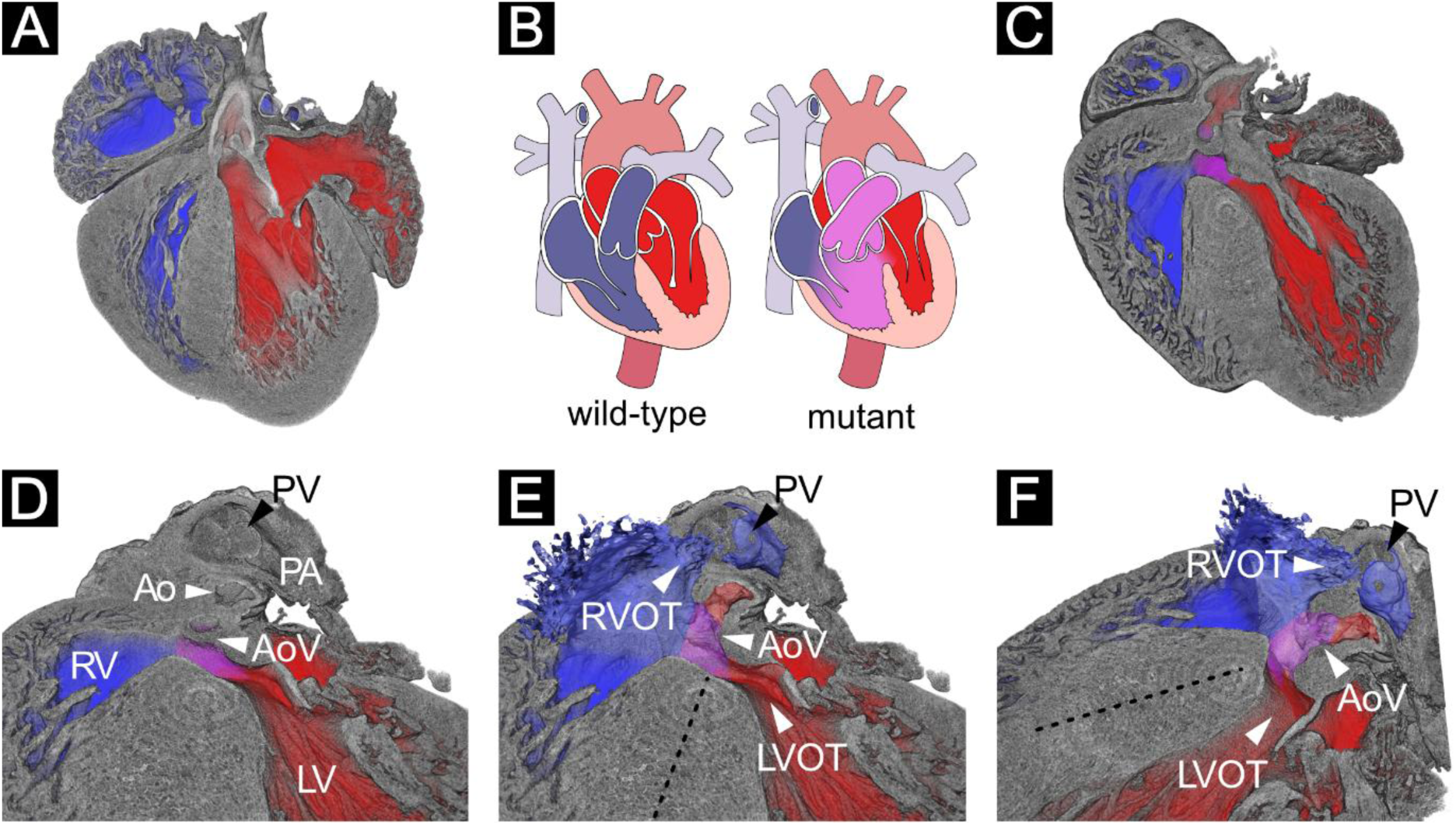
Example of application of PC-µCT to cardiac phenotyping. **A –** Volume rendering of a wild-type E18.5 mouse heart, digitally cropped to show a 5-chamber view. The luminal volumes of the cardiac chambers are color-coded to distinguish laterality, right-sided in blue and left-sided in red. **B** – Schematic drawings of a wild-type heart and of the defect observed in the mutant heart. **C** – Volume rendering of the heart of an *Adamts6*-deficient E18.5 mouse, in a view matching the wild-type in A. Between right and left side, a ventricular septal defect is clearly visible (in purple). **D** – Close-up view of the ventricular septal defect in C, with an additional part of the rendering extended in the vertical direction, to show the pulmonary artery as well. The overriding of the aorta defines a double outlet right ventricle type of defect (DORV). RV: right ventricle, LV: left ventricle, Ao: aorta, AoV: aortic valve, PA: pulmonary artery, PV: pulmonary valve. **E** – As in D, but adding semi-transparent surface mesh renderings of part of the luminal volumes, to better show right ventricular outflow tract (RVOT), left ventricular outflow tract (LVOT) and septal defect connection. A dashed line is added to the middle of the septum, confirming the overriding aorta. **F** – As in E, but from a different view that better shows the septum and the overriding.

While the ventricular septal defect is visible already from the sectioning plane in Figure 3-C, by leveraging the volumetric information provided by PC-µCT (Figure 3-D-E-F) it is possible to observe that both aorta and pulmonary artery arise predominantly from the right ventricle, defining a double outlet right ventricle type of defect. In Figure 3-D, a close-up of Figure 3-C shows the ventricular septal defect, with aorta and aortic valve above it; the volume rendering is also extended in the vertical direction, to show pulmonary artery and pulmonary valve. In Figure 3-E, the right ventricular outflow tract is added as a semi-transparent mesh surface rendering, to clarify the opening of the right ventricle. The septum is also indicated with a dashed line, to strengthen the argument of the defect being a double outlet right ventricle. Figure 3-F offers a different view. Contrasting with histological assessment from a single section, which was limited by the choice of initial block orientation, the present analysis eliminated ambiguity of detection and enabled precise delineation of this complex defect.

### 3.2. Integration of histology and fluorescent staining with PC-µCT

After PC-µCT, the paraffin-embedded samples were sectioned and stained. The histology and fluorescent staining images were then integrated back into the CT data.

In between the performing of PC-µCT and the scanning of the histological sections, the sections undergo a series of transformations (sectioning, de-paraffinization, re-hydration, staining, de-hydration) that inevitably deform the tissue. In order to register (i.e. digitally combine) the two different datasets with high precision, a data processing pipeline consisting of multiple steps was developed. The pipeline is summarized in Figure 4 and described in detail in the Methods section.

**Figure 4.**
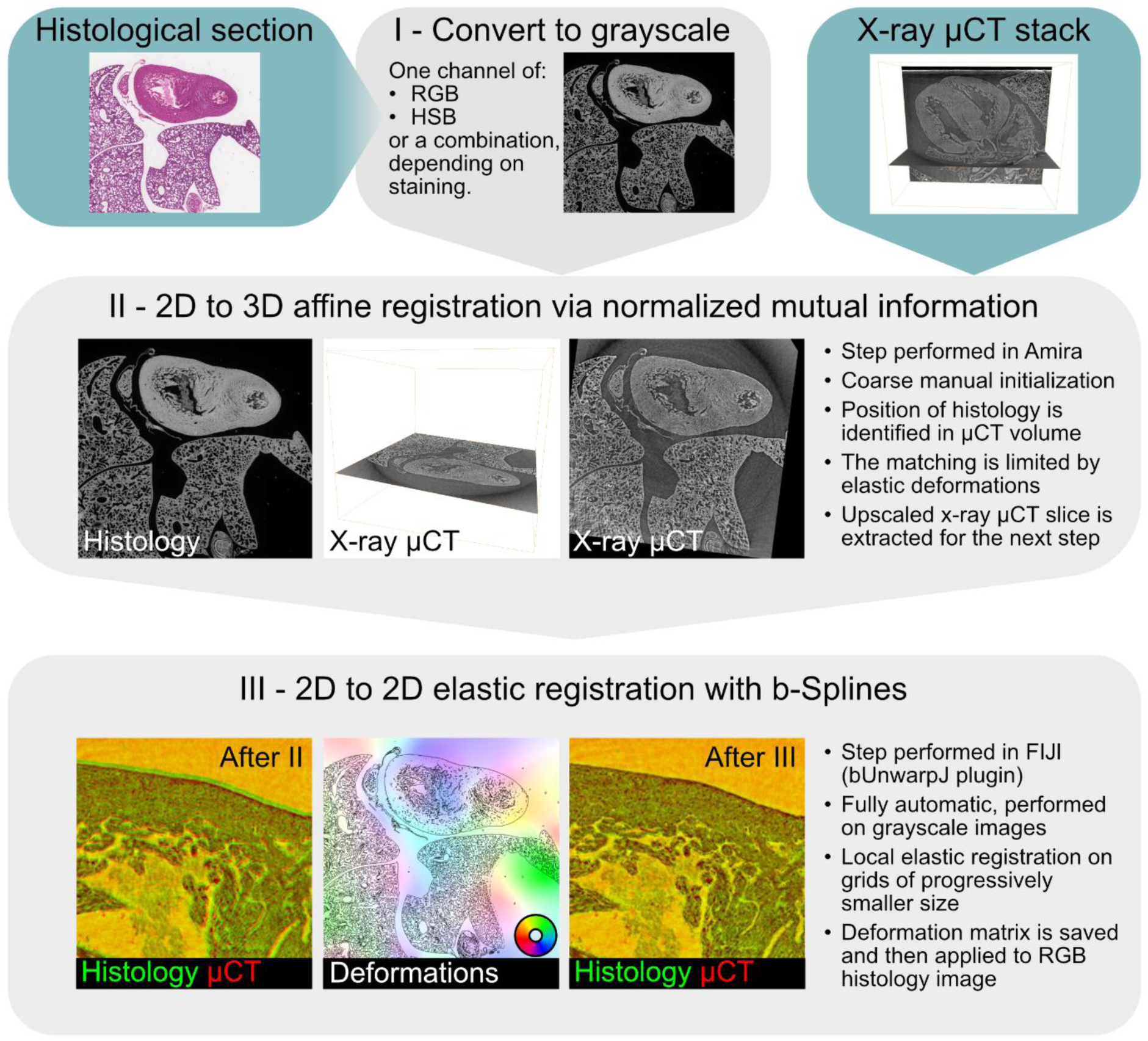
Schematic overview of the data registration pipeline used in this work. The input data, histological section and x-ray µCT stack, are shown with a cyan background. From top to bottom, the pipeline is then formed by three main steps (gray background). **Step I** is a grayscale conversion of the histology data, while trying to match the feature contrast provided by the x-rays; to this end, one channel from the RGB (red, blue, green) or HSB (hue, saturation, brightness) color decomposition is typically used, depending on the staining. The grayscale histology and the µCT stack are then loaded in Amira (Thermo Fisher Scientific) for **step II**, a 2D-to-3D affine registration. This procedure is semi-automatic, requiring a first coarse manual positioning of the histology into the µCT volume, followed by an automatic refinement using a normalized mutual information score. At this point of the pipeline, the match between histology and PC-µCT typically looks acceptable at low magnification but is far from a 1-to-1 match when evaluated at a zoomed-in level. This is due to local deformations of the tissue, which cannot be fixed with the global transformations performed so far. This is addressed in **step III**, where the identified matching slice in the µCT volume is upscaled to the histology resolution and the two 2D images are then processed for 2D-to-2D elastic registration using the bUnwarp plugin in FIJI. The plugin is fully automatic and performs local elastic registration using b-splines estimated on grids of progressively smaller size. The green (histology) and red (µCT) overlapping images show the improvement introduced by step III compared to only step II. A map of the estimated deformations (hue shows the direction, saturation the relative intensity) is included to show the highly localized character of these changes. All the mathematical transformations applied to the grayscale-converted histology images can finally be applied to the original images, which can then be loaded in Amira to produce the other figures in this section.

An example of the result for hematoxylin and eosin (H&E) staining is provided in Figure 5, in which three H&E-stained sections were identified in the PC-µCT scan presented in Figure 1. Here they are shown combined with a volume rendering with carved corner (Figure 5-A) or with mesh surface renderings of heart and coronary circulation (Figure 5-B, with inset showing the good match between histology and segmentation of the coronaries). A direct comparison between the same slice in SRPC-µCT and histology after the registration pipeline is provided in Figure 5-C and -D, respectively.

**Figure 5.**
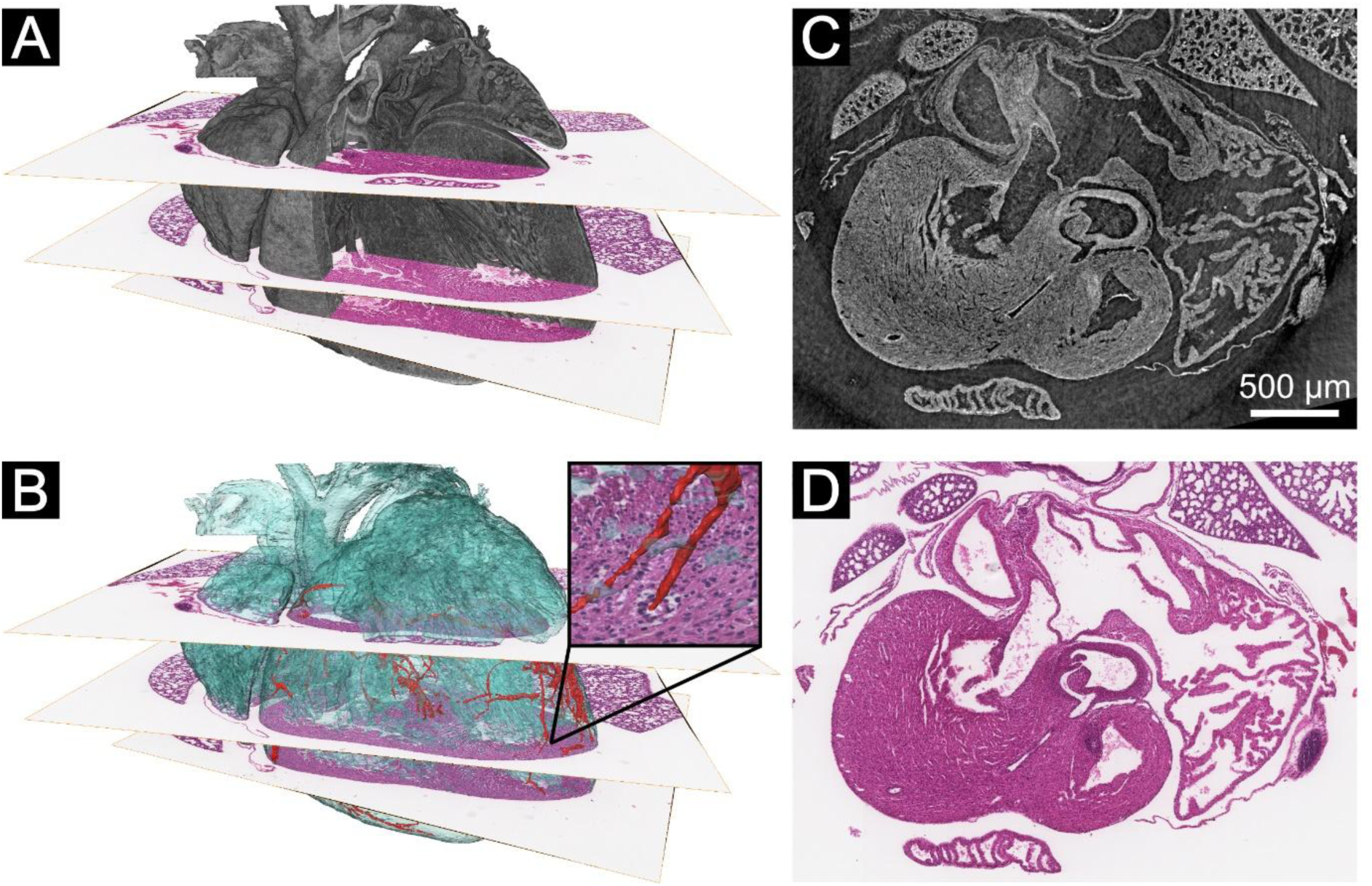
Integration of standard hematoxylin and eosin (H&E) staining with PC-µCT. **A –** Three different H&E-stained sections are integrated into the volume segmented in Figure 1. A corner of the volume rendering is carved to show more of the H&E-stained sections. **B** – Using the surface renderings of Figure 1-D, the match between histology and µCT is evident down to the registration of coronary arteries (inset). **C** – The SRPC-µCT digital slice matching one of the histology images. **D** – The H&E image matching C.

The registration pipeline was also tested with fluorescent staining (FS) data; the results are exemplified in Figure 6. Figure 6-A shows how the semilunar valves (arrows) with connections to the aorta and the main pulmonary artery could be visualized with a combination of PC-µCT slices and volume rendering (in pink, to more easily distinguish it from the grayscale 2D slices). In Figure 6-B, a tissue section stained with immunofluorescence for α-actin, to visualize smooth muscle cells, and DAPI for cell nuclei, has been integrated within the volume. The smooth muscle cells in the walls of the great arteries (aorta and pulmonary artery), as well as in a coronary artery (arrow) are clearly positive for α-actin, while DAPI provides an overview of tissue cellularity. Figure 6-C shows only the slices, without the rendering, to make the α-actin in the arterial walls more visible and also highlight how volumetric information is important to correctly understand the morphology. Figure 6-D, -E, -F combine a volume rendering based on the PC-µCT data with FS for hyaluronan binding protein, which was localized into valve leaflets as expected. In E and F, to better show this localization, the pulmonary valve and trunk were added as a semi-transparent mesh surface rendering.

**Figure 6.**
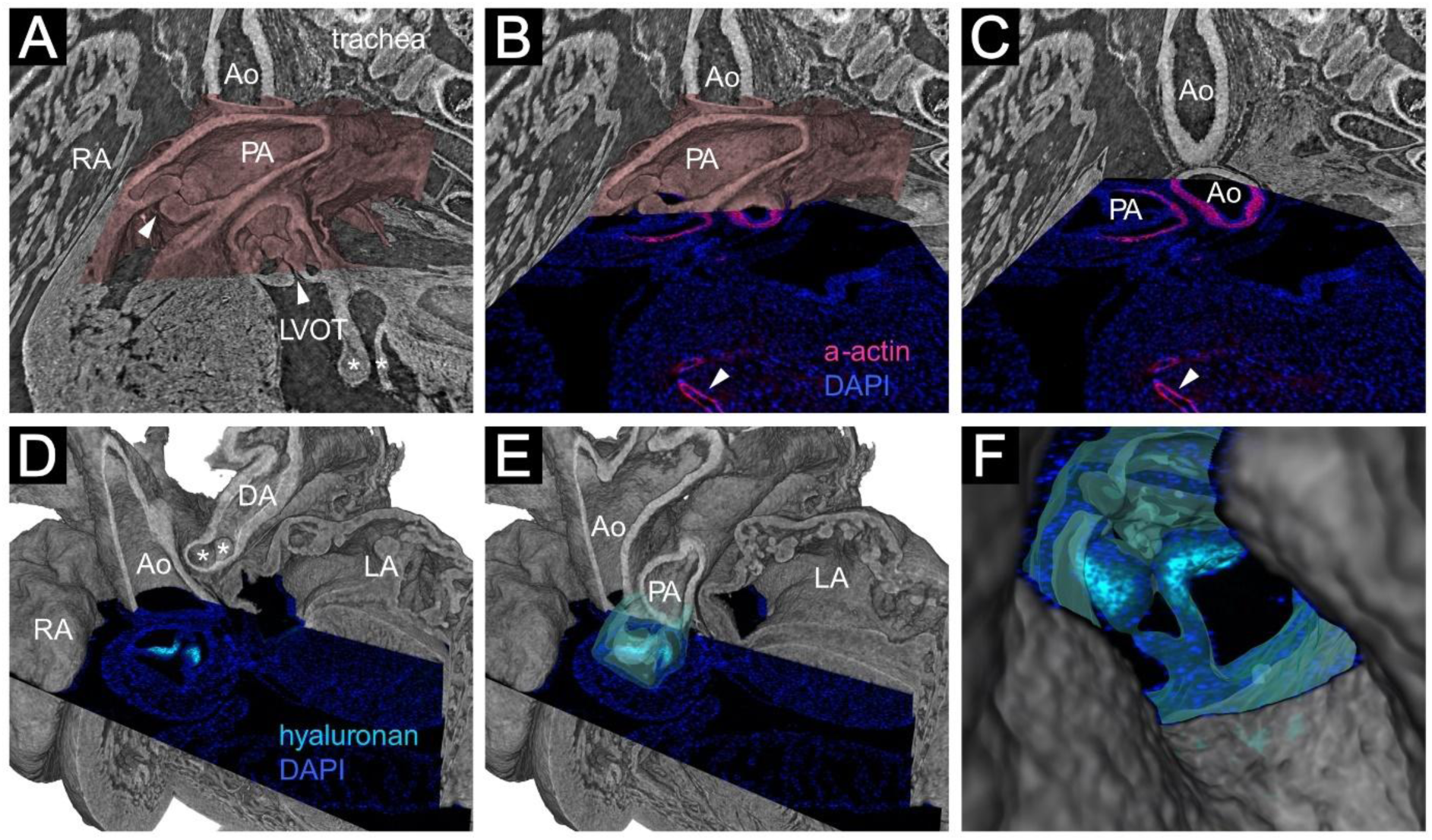
Integration of molecular staining with PC-µCT. **A** – Different µCT slicing planes (grayscale), combined with a volume rendering (in pink, to clearly separate it from the slicing planes), can be used to identify several morphological features, such as the semilunar valves (arrowheads) connecting to aorta (Ao) and pulmonary artery (PA), and the mitral valve (asterisks). RA: right atrium, LVOT: left ventricular outflow tract. **B** – An immunofluorescence section stained for α-actin (in red), to visualize smooth muscle cells, and DAPI (in blue), for cell nuclei, is integrated into the volume. The walls of Ao, PA, and of a coronary artery (arrowhead) are clearly positive for α-actin. **C** – The volume rendering is removed to make the α-actin in the Ao more visible, and to show the importance of the volumetric information to correctly understand the morphology. **D** – Different view of the same heart (now rendering in grayscale, since no slicing planes are used), combined with a section stained for hyaluronan (in cyan) and DAPI. Hyaluronan is highly localized in the valve leaflets, as expected. Several morphological features are recognizable, like the ductus arteriosus (DA) and the branching of the pulmonary artery (asterisks). LA: left atrium. **E** – Like D, but adding the pulmonary valve and trunk as semi-transparent mesh surface rendering and adjusting the cropping of the volume rendering. **F** – Like E, different point of view (view coming from the DA inside the PA), to highlight the registration between the datasets.

### 3.3. Integration of spatial transcriptomics with PC-µCT

The same pipeline was successfully applied to spatial transcriptomics data (Xenium platform) as well, the results of which are shown in Figure 7.

**Figure 7.**
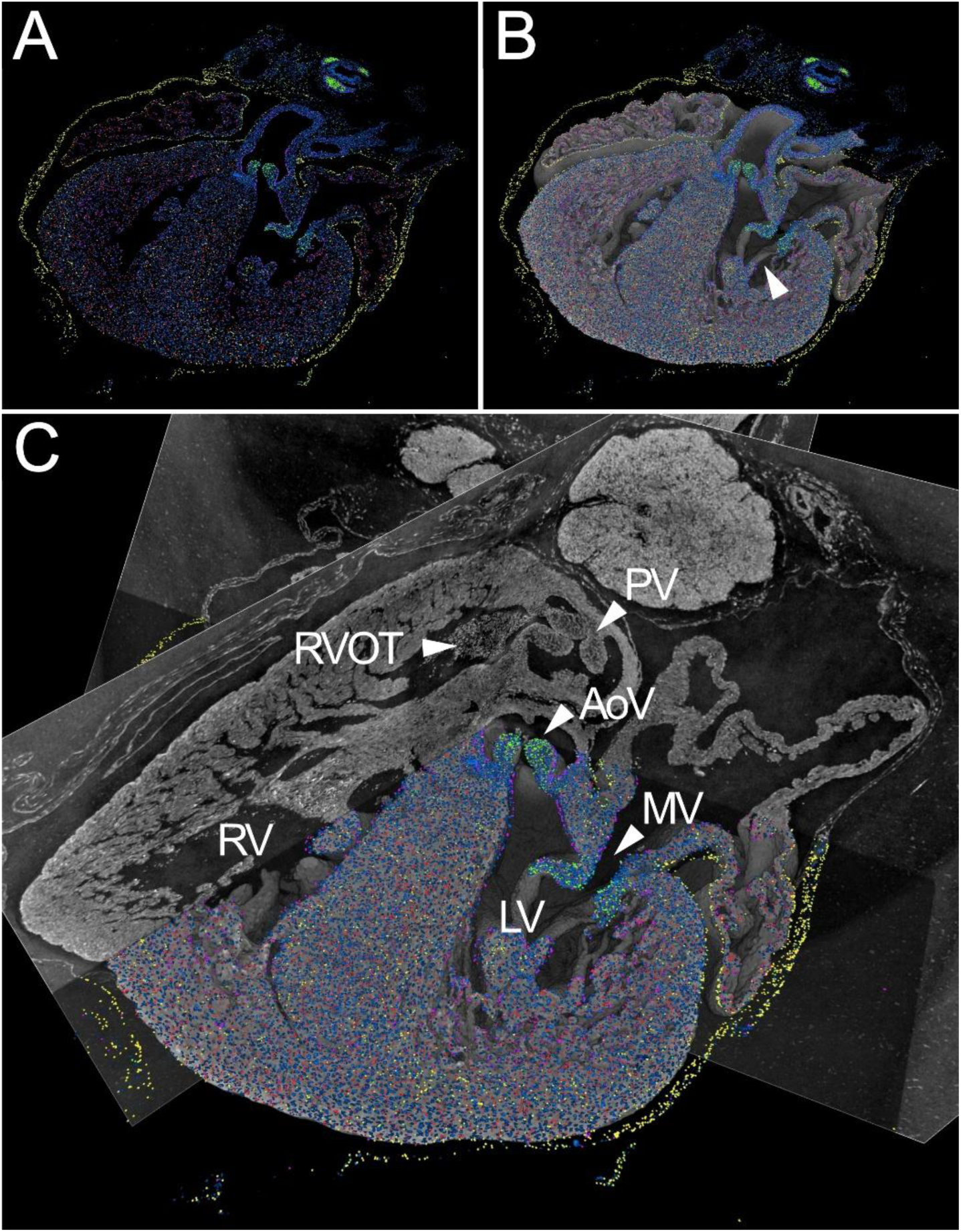
Integration of *in situ* sequencing (ISS) data with PC-µCT. **A** – From the ISS data (Xenium platform), four genes with distinct role and spatial localization in the heart were selected and displayed on top of DAPI (blue). The genes are: *Tnni3k* (red), localized in the myocardium; *Wt1* (yellow), in the epicardium; *Foxc1* (purple), in the endocardium; *Hapln1* (green), in the valves and cartilage. **B** – The ISS slice is overlapped to the matching PC-µCT volume rendering. Morphological information, such as the papillary muscles (arrowhead) connecting to the mitral valve, are readily visible. **C** – Combining rendering and off-plane digital sections provides greater structural context. RV: right ventricle, LV: left ventricle, RVOT: right ventricular outflow tract, AoV: aortic valve, PV: pulmonary valve, MV: mitral valve.

In Figure 7-A, four transcripts, of genes known for distinct roles and spatial localization in the heart ^35,36^, were selected from among the 5000 total genes in the Xenium data and displayed in four distinct colors, on top of DAPI (in blue). Troponin I-interacting kinase (*Tnni3k*, in red) regulates cardiomyocyte growth, contractility and conduction system development, and was localized prevalently in the myocardium; Wilms tumor 1 (Wt1, in yellow) is essential for epicardium formation and was predominantly localized there; Forkhead box C1 (*Foxc1*, in purple) is involved in cardiac morphogenesis, and was previously reported to be expressed mainly in the endocardium, which was confirmed here; Hyaluronan and proteoglycan link protein 1 (*Hapln1*, in green) supports structural integrity of the extracellular matrix. As expected, based on previous publications and our FS with hyaluronan binding protein shown in Figure 6, *Hapln1* expression was mainly seen in cardiac valves (as well as in tracheal cartilage).

In Figure 7-B, spatial transcriptomics data is combined with the matching volume rendering obtained from PC-µCT, providing 3D context. As an example, the mitral valve papillary muscles are clearly identifiable (arrowhead), connecting to the leaflets. Figure 7-C also adds to the rendering virtual slices out of the spatial transcriptomics plane, showing the position of the pulmonary valve and the right ventricular outflow tract.

After spatial transcriptomics, the section was also stained with FS for CD31 (to label endothelial cells) and for hyaluronan. It was then stained with H&E as well. All these results were registered to the PC-µCT data with the same pipeline. Figure 8 shows how PC-µCT, spatial transcriptomics, FS and H&E data can all be combined in the same view. Here, the visualized *in situ* sequencing (ISS) data corresponds to one cell cluster, associated to a valvular subpopulation of extracellular-matrix-producing fibroblasts (in green, on top of DAPI in blue). In addition to the cluster, two selected transcripts associated with hyaluronan, hyaluronan synthase 2 (*Has2*) and *Hapln1*, are explicitly shown (in pink and yellow respectively). As expected, they are clearly localized to the leaflets of the aortic and mitral valves, just like the hyaluronan FS (in cyan). FS for CD31, to visualize endothelial cells, is shown in red.

**Figure 8.**
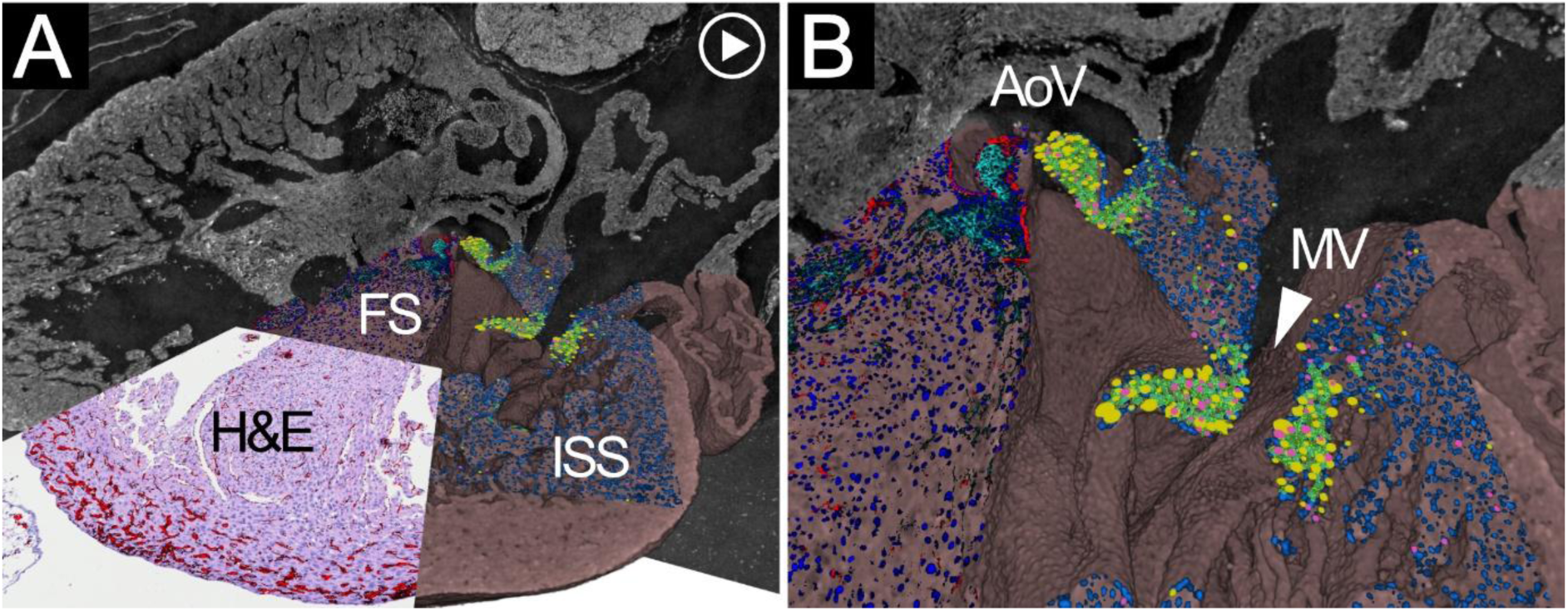
Integration of PC-µCT with standard histology, fluorescent staining (FS) and spatial transcriptomics (*in situ* sequencing, ISS). **A** – The ISS slice shown in Figure 7 was subsequently stained with FS for DAPI (in blue), hyaluronan (in cyan) and CD31 (in red), and later re-stained with H&E. All the images were registered together and are shown in separate quadrants, overlapped to the corresponding PC-µCT volume rendering (tinted in pink to distinguish it from the virtual slices) and combined with virtual slices to provide additional volumetric context. The ISS data here shown correspond to a single cell cluster, associated to a valvular subpopulation of extracellular-matrix-producing fibroblasts (in green, on top of DAPI in blue). From this cluster, two selected transcripts associated to hyaluronan (*Has2* and *Hapln1*) are explicitly shown (in pink and yellow respectively). In the H&E quadrant, FS for CD31 is also overlapped, to visualize the endothelial cells. **B** – Close-up view on the aortic valve (AoV) and the mitral valve (MV), showing smaller details of the FS and ISS data. The selected ISS cluster, and the two selected transcripts associated to hyaluronan, are highly localized in the valve leaflets, just like the hyaluronan FS in cyan, as expected. **Video-clip 2** illustrates how these combined views were created.

Video-clip 2 illustrates how these combined views were created and provides a better spatial understanding.

### 3.4. Laboratory-based PC-µCT

After SRPC-µCT, one sample was also imaged on two LBPC-µCT scanners, a commercial one (Exciscope Polaris) and a high-resolution custom-made one ^37^.

Prior to the scans, excess paraffin was manually removed from around the sample with a scalpel, as detailed in the Methods section.

The commercial system was used in a configuration which provided a pixel size somewhat comparable to the synchrotron data (2.46 µm vs 1.63 µm). Five different total scan times were tested, from 5 minutes (to directly compare with the synchrotron results) to 8 hours. The results are summarized in Figure 9.

**Figure 9.**
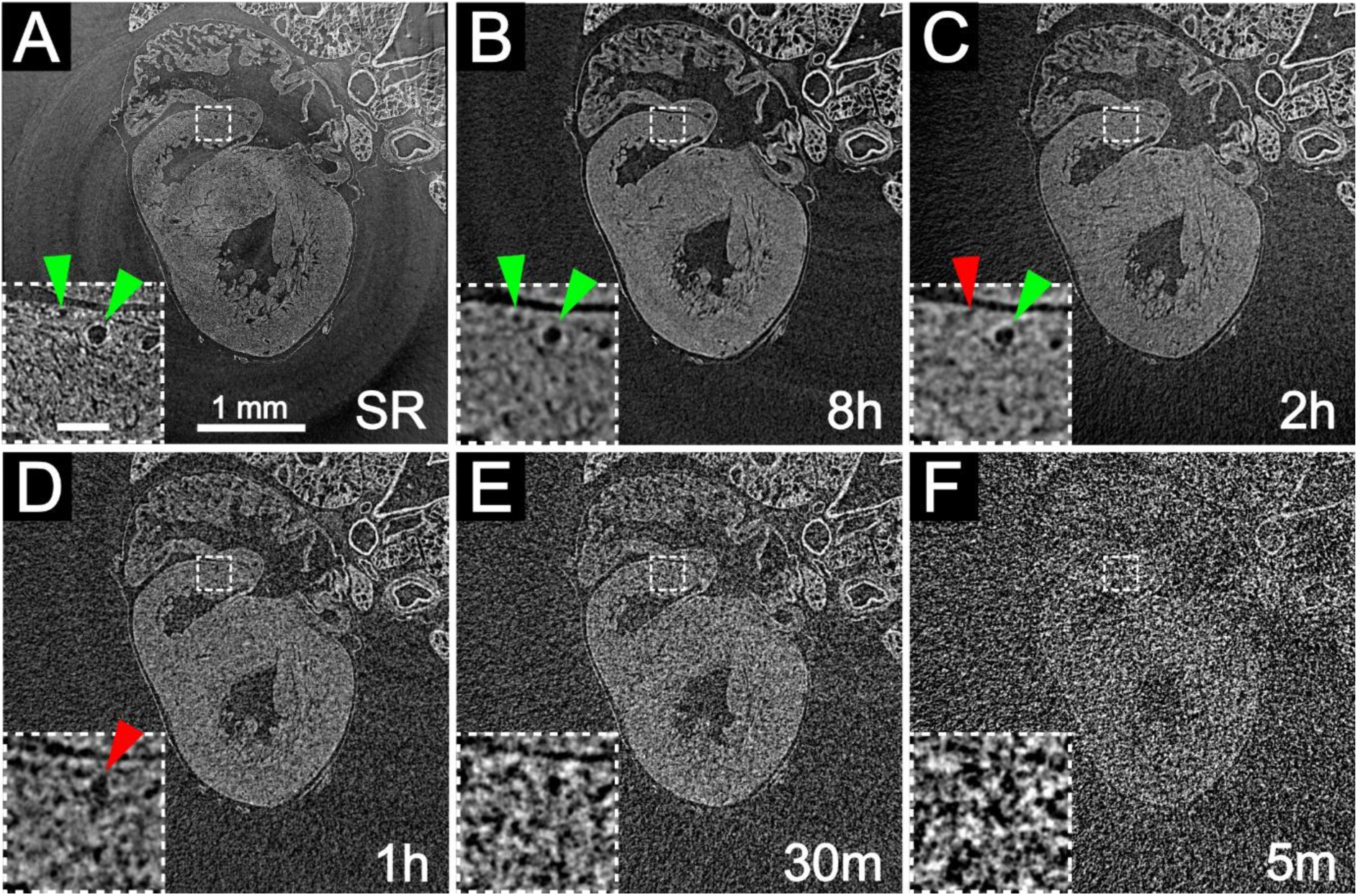
Qualitative comparison between a 5-minutes-long SRPC-µCT scan (panel **A**) and LBPC-µCT scans with different total scan times (8 hours, 2 hours, 1 hour, 30 minutes and 5 minutes in panels **B-to-F** respectively) performed on a commercial system (Exciscope). The field-of-view of the LB systems is cropped to the one of the SR data (full field-of-view is shown in Supplemental Material). In each figure, the inset in the bottom left corner is a magnified view of the same region of interest (dashed outline). In the inset, arrowheads point to coronary arteries of different size. The arrowheads are color-coded to indicate in which scan they can be seen (green) or when they start to be lost (red). Scalebar in the inset: 100 µm.

In the figure, all the datasets have been registered to each other, and a region of interest matching the field-of-view of the SRPC-µCT scan has been extracted from the LBPC-µCT data. The field-of-view provided by the laboratory setup was significantly larger than shown (approximately twice as large in each direction, owing to a larger detector size compared to the synchrotron setup). Most of this was unused for the heart scan, but it would allow for the scanning of larger samples without having to compromise the effective pixel size. An image of the full field-of-view is provided in Figure S1, Supplemental Material.

From a qualitative evaluation of Figure 9, it is apparent that, if only the larger structure of the heart is of interest, for example to identify ventricular septal defects as in Figure 3, a 2-hours-long LBPC-µCT scan might be sufficient. Even 1 hour or 30 minutes might be acceptable, depending on the size of the defect and on the noise requirements for the segmentation.

Smaller details, like the coronary circulation, would require longer scan times, as shown in the insets, which correspond to magnified views of the dashed region-of-interest in the scans. Two arrowheads point to a larger coronary vessel (on the right) and a smaller one (on the left); the arrowheads are color-coded, with green indicating that the coronary vessel is visible, and red meaning that the details of the coronary vessel start being lost.

Details that are even smaller than the coronary vessels, such as individual cardiomyocytes, which are distinguishable in the SRPC-µCT dataset, are below the resolution limit of the LBPC-µCT system. While not strictly essential for the applications described in this work, they would be crucial for studies in which cardiomyocyte orientation, extracted for example with structure tensor analysis ^12,15^, would be of interest. Higher resolution could in theory also improve registration accuracy, by providing more detailed information about the underlying structures, thereby better constraining the alignment between datasets.

The custom-made laboratory scanner provided similar results to the commercial one (shown in Figure S2, Supplemental Material). This system was initially included because of its potential to reach even higher resolutions (< 1 µm) ^37^, but the relatively large sample size, especially due to the paraffin still surrounding the heart, impaired the available magnification. Nevertheless, it still managed to achieve results comparable to the commercial scanner when used with lower magnification settings.

## 4. Discussion

This paper, to our knowledge, shows for the first time how high-resolution 3D imaging, in the form of SRPC-µCT, could be successfully combined with ISS-based spatial transcriptomics, fluorescent staining and conventional histology. All 2D methods integrated into the CT volume could be performed sequentially on a single tissue slice. The methodological pipeline is described in detail, and this multimodal approach will be of great value not only for efficient murine cardiac phenotyping, but also for combining structural and molecular information in many different fields of research.

Rapid and accurate phenotyping of mutant mouse strains will be of value not only for the research groups who now use more time consuming or destructive methods, but also has potential to reduce the number of animals needed for confirming a phenotype, in agreement with the 3R principle to replace, reduce and refine the use of animals for research ^38^ (as cited in ^39^).

The possibility to virtually section the tissue in standard echocardiography views ^31^ will facilitate communication with physicians working clinically with congenital heart disease, who may be unfamiliar with PC-µCT. For this study, the work to find the correct imaging planes was done manually by adjusting the slicing planes in Amira. However, one could envision a future method where landmarks are added to certain points of interest, such as features within the valves and the apex, through which the virtual imaging planes could be automatically fitted. Such landmarks could also potentially be used to train artificial intelligence (AI) or machine learning approaches aimed at fully or partially automated cardiac phenotyping in the future.

Importantly, no specific adjustments of the protocols previously used for preparation of hearts or for SRPC-µCT were needed for this project. Standard tissue fixation and paraffin embedding were used, taking care to remove as much blood as possible to make segmentation easier, combined with careful avoidance of air bubbles in the paraffin blocks to ensure optimal image quality. Before performing the Xenium experiment, a comparison of RNA quality ^40^ was made using sections from a heart which had not been imaged with X-rays, and sections from a heart imaged twice with standard protocols for SRPC-µCT (at the ANATOMIX beamline ^41^ of the SOLEIL synchrotron in Paris, France, and at the DanMAX beamline ^42^ of the MAX IV synchrotron in Lund, Sweden) and once at a laboratory-based micro-CT scanner (Exciscope Polaris) at the Technical University of Denmark in Copenhagen (DTU), also using standard protocol imaging for this type of soft tissue. No obvious differences in detection of common RNAs were observed between the two hearts, indicating that standard protocols for tissue processing and µCT can be used when planning for subsequent ISS. This confirms previous findings regarding preserved RNA integrity following SRPC-µCT ^4^.

The laboratory scans showed that, if measurement speed is not a concern, laboratory-based solutions could be a more accessible alternative to synchrotrons, depending on the length-scale of the features of interest. The longest tested scan time on the commercial scanner could resolve microanatomical features as small as coronary vessels with a cross-section of ∼10 µm, although these were quickly lost with shorter scan times. If larger defects are of interest, we showed how scan times down to 1 hour or possibly even 30 minutes per sample could still provide satisfactory results. It is important to mention that the comparison was not entirely fair, as the synchrotron scan was performed with the heart embedded in a standard paraffin block with a size of roughly 3 x 2 x 1 cm^3^, while prior to the laboratory scans the block was carved into a ∼1-cm-diameter cylinder centered around the heart, removing the excess paraffin. If the sample had been processed in a similar way, we could have expected even shorter scan times at the synchrotron, as well as a probable reduction of ring artefacts and local tomography artefacts. Nevertheless, laboratory scanners show great potential if they can be accompanied by an ad-hoc standardization of the embedding protocols. This was even more relevant for the custom made high-resolution system, which here showed similar performance to the commercial scanner but could in principle offer even higher resolution if more paraffin could be removed from around the sample.

Having a 3D visualization of a tissue sample prior to sectioning is useful for selecting regions of interest for subsequent multimodal analysis. Imaging the tissue already embedded in a cassette which will fit the microtome used for sectioning, in addition to keeping the block as horizontal as possible while performing the CT, is particularly helpful. There will always be tissue distortion when re-hydration and de-hydration of tissue is performed, but it is preferable if tissue movement and/or detachment can be minimized by handling the tissue with care, due to the limit to how much distortion the pipeline can compensate.

A single tissue section was used to combine the different methods and we show only one of many cell clusters combined with transcripts actively expressed in this specific cluster to illustrate the visualization possibilities. However, we have previously developed virtual histological staining (Elastin van Gieson and H&E) by using serial sectioning and staining of every 10^th^ section for generative adversarial network training purposes (manuscript currently in second round of revision in the Journal of the Royal Society Interface). A similar approach could possibly be used for AI-assisted identification of cell clusters within CT volumes.

In summary, this study shows how high-resolution 3D imaging (SRPC-µCT) was successfully combined with spatial transcriptomics, fluorescent staining and conventional histology on a single tissue slice. The approach described herein has the potential to be a useful proof-of-concept guide for additional studies that will encompass a multidisciplinary, multimodal approach.

## 5. Methods

### 5.1. Ethics statement

The samples were obtained from Lund, Sweden and Cleveland, Ohio, USA. All animal experiments from Sweden were approved by the Malmö/Lund (Sweden) Ethical Committee for Animal Research (ethical permit numbers M173–19 and M2780–24), in accordance with guidelines from the Swedish Board of Agriculture. All experiments were performed in line with the laws and regulations issued by the Swedish Board of Agriculture. Experiments involving live animals were reported according to the ARRIVE guidelines. The animal experiments obtained from the USA were approved by the Cleveland Clinic Institutional Animal Care and Use Committee (protocol 2018–2450).

### 5.2. Tissue preparation

The hearts shown in Figures 1-2–5–6–9 were healthy control embryonic hearts imaged for a phenotyping study of laminin γ1 conditional knockout mice ^11^. Animals were maintained at the Biomedical Center animal facility (Lund, Sweden), where routine pathogen surveillance was performed. Genotyping was carried out according to Jackson Laboratory protocols. Timed matings were established to obtain embryos for analysis. Pregnant females were sacrificed at day 18.5 of embryo gestation and embryos were collected. Embryos were decapitated and positioned supine for fixation, and a tail sample was collected for genotyping. The peritoneal cavity was opened by a midline incision extending to the trachea. The trachea was perforated using a blunt G30 needle and lungs were inflated with Histochoice (Sigma). The heart–lung complex was then isolated and fixed in Histochoice overnight at 4°C.

Histochoice-fixed tissues were dehydrated and paraffin embedded. Samples were transferred to PBS overnight at 4°C and then sequentially through 45%, 70%, 95%, and 100% ethanol followed by xylene (approximately 1 h per step at room temperature). Tissues were then incubated in paraffin at 60°C (two changes, 1.5 h each) and embedded.

The hearts presented in Figures 3-7–8 were from E16.5 wild-type and from an *Adamts6*-deficient strain. *Adamts6* mutant mice were previously described ^32,33^ and maintained on a C57BL/6 J background. Age-matched wild-type C57BL/6 embryos were used as controls. Mouse embryos were generated by controlled matings, using the observation of a vaginal mucus plug to designate E0.5. The hearts were fixed in 4% paraformaldehyde, processed and embedded in paraffin.

### 5.3. Synchrotron-based phase-contrast µCT

The samples presented in Figures 1-6 and Figure 9 were scanned with SRPC-µCT at the X02DA TOMCAT beamline ^43^ of the Swiss Light Source (SLS, Paul Scherrer Institute, Villigen, Switzerland).

PC-µCT was performed using the free space propagation method, which does not require any additional equipment aside from a coherent X-ray source and a detecting system placed at a specific propagation distance. The X-ray beam, produced by a 2.9 T super-bending magnet inserted in the 2.4 GeV storage ring, was monochromatized to an energy of 21 keV by using a double-multilayer monochromator. The detecting system consisted in a 20-µm-thick LuAG:Ce scintillator coupled to a 4X magnifying objective (Optic Peter) and a PCO Edge sCMOS detector, resulting in a field-of-view of 4.2 x 3.5 mm^2^ with an effective pixel size of 1.63 x 1.63 µm^2^. The sample was placed on a rotating stage in-between source and detector, with a sample-to-detector propagation distance of 19 cm. The acquisition time was set to 80 ms per projection, for a total exposure time of about 2.4 min for a full scan (180° rotation) with 1801 projections. Including 100 flat images and 30 darks, plus motor overhead, the total scan time was about 4.3 min.

Following acquisition, the projections were processed for phase retrieval using Paganin’s method ^44^ (δ = 3.7e-8, β =1.7e-10). Tomographic reconstruction was then performed using the gridrec algorithm ^45^. The obtained data were digital volumes of 2560 x 2560 x 2160 pixels (corresponding to a physical volume of 4.2 x 4.2 x 3.5 mm^3^), with a 16-bit pixel depth. The sample used for Figures 7-8 was imaged at the ANATOMIX beamline ^41^ of the SOLEIL synchrotron. The x-rays were produced by an U18 cryogenic in-vacuum undulator, which was set to a gap of 5.5 mm. The beam was filtered to a pink beam using a combination of filters (26 um Au, 0.1 mm Cu) that corresponded to a mean energy of approximately 30 keV (25 keV peak). The detecting system consisted in a 20-µm-thick LuAG:Ce scintillator coupled to a 5X magnifying objective (Mitutoyo) and an ORCA Flash 4.0 V2 sCMOS detector (Hamamatsu), for an effective pixel size of 1.3 µm and a 2.7 x 2.7 mm^2^ field-of-view. The sample-to-detector distance was set to 20 cm. A full scan (180° rotation) consisted of 2000 projections with a 100 ms acquisition time each, for a total exposure time of about 3.3 min. Considering flat and dark images, and motor overhead, the total scan time was around 5 min.

Following acquisition, the data reconstruction pipeline was performed using the beamline routines, based on PyHST2 ^46^. The phase information was retrieved with Paganin’s method ^44^ (with a Paganin length of 130 µm; this parameter is described in the PyHST2 documentation ^47^ and for a 30-keV mean photon energy and a 20-cm sample-to-detector distance corresponds to a δ/β of around 650). An unsharp mask (filter length 2, filter coefficient 0.3) was also applied to reduce the excessive smoothing introduced by the phase retrieval method. Ring removal was performed using the default ring correction in PyHST2, described in ^46^ and named “double flat field correction” in the PyHST2 documentation ^47^. The volumes were then reconstructed using filtered back projection. The obtained data were digital volumes of 2048 x 2048 x 2048 pixels (corresponding to a physical volume of 2.7 x 2.7 x 2.7 mm^3^), with a 16-bit pixel depth.

### 5.4. Laboratory-based µCT

After SRPC-µCT, one of the samples scanned at TOMCAT was imaged with two different laboratory setups. Before performing LBPC-µCT, excess paraffin was manually removed from around the sample with a scalpel, obtaining a paraffin cylinder of approximately 1 cm diameter with the sample roughly in the center. This step was quite important for the laboratory setups, both for time concerns (less paraffin absorbing the X-rays means shorter scan times) and for geometrical requirements of the custom-made laboratory setup (in which, to achieve higher geometrical magnification, it was necessary to position the sample as close to the source as possible).

The first system was a version of the commercial scanner Exciscope Polaris, equipped with an Excillum MetalJet D2+ 70 kV source (with the I1 target alloy) and a Photonic Science Gsense XL CMOS camera (4096 x 4096 pixels, 13.1 µm pixel size). The source was run at 187 W of power (average energy 17 keV), with an e-beam spot size of 10 x 60 μm^2^. The scanning geometry was set to have a 150 mm source-to-object distance and an 800 mm source-to-detector distance, resulting in a 5.3x cone-beam magnification, corresponding to an effective pixel size of 2.46 µm and an effective propagation distance of 122 mm. A total of 3601 projections were acquired, uniformly covering a 360° rotation of the sample. Five different scans were performed, with different exposure times per projection (0.083 s, 0.5 s, 1 s, 2 s and 8 s). This resulted in total scan times ranging from 5 min (comparable to the SRPC-µCT one) to 8 h. The acquired images were processed for artefact reduction (Exciscope proprietary algorithm), phase retrieval (using Paganin’s method ^44^) and tomographic reconstruction (filtered back projection for cone-beam geometry using the commercial software MITOS).

The second system was a custom-made setup ^37^ comprising an Excillum NanoTube N2 60 kV source and a DECTRIS EIGER2 R 500K photon-counting detector (1028 x 514 pixels, 75 µm pixel size). The source was run at 2.4 W (average energy 16 keV), with a 1.2 µm e-beam spot size. The source-to-object distance was set to 16.16 mm and the source-to-detector distance was set to 207.20 mm, corresponding to an effective pixel size of approximately 5.8 µm. One tomographic scan was performed, with 1200 projections and 15 s exposure time, for a total scan duration (including 30 flat field images and 36×2 pre- and post-alignment images, plus motor overhead) of 5.5 hours. The system is optimized for higher resolutions using much smaller samples, but the relatively large sample size imposed restrictions on the available magnification while keeping reasonable scan times. The acquired data were processed with a custom-developed pipeline written in Python, detailed in ^37^. As part of the pipeline, phase-retrieval was performed with the method by Paganin et al. ^44^ (δ = 3.7e-8, β =1.7e-10). Tomographic reconstruction was carried out with a GPU implementation of the Feldkamp-Davis-Kress algorithm ^48^ in the ASTRA toolbox ^49^. In the reconstruction step, a super resolution factor of 2 was selected ^50^, essentially interpolating the reconstruction onto a 2x finer grid, for a final effective pixel size of 2.9 µm.

### 5.5. Histology

For the H&E staining, the paraffin-embedded samples were sectioned into 5-μm-thick sections. The sections were deparaffinized in xylene and rehydrated through a series of graded alcohols. H&E staining was performed according to a standard protocol. Sections were scanned using Aperio’s Scanscope CS2 (with Scanscope console v.8.2.0.1263) at 20x magnification. Images were created using Scanscope software (Aperio).

For the IF staining, 3-μm-thick sections were deparaffinized and rehydrated through decreasing concentrations of alcohol. Heat-induced antigen retrieval (HIER) was performed using the DIVA decloaker (Biocare Medical) for 5 minutes at 110°C in a decloaking chamber (Biocare Medical). After a quick wash, Background Punisher (Biocare Medical) was applied for 10 minutes at room temperature (RT) for 10 minutes to block nonspecific binding. Consequently, tissues were incubated overnight at 4°C with biotin-conjugated hyaluronic acid binding protein (HABP) (1:100, AMS_HKD_BC41, Amsbio). Cy5-conjugated streptavidin (1:200, SA1011, Invitrogen) was used for detection. Smooth muscle actin (SMA) and nuclei were stained with a Cy3-conjugated primary SMA antibody (1:200, C6198, Sigma Aldrich) and DAPI (1:1000, MBD0015, Sigma Aldrich), respectively, for 1 hour at RT, along with the streptavidin incubation. 1% BSA in PBS was used as the antibody diluent, and slides were sealed with DAKO fluorescence mounting media (S3023, Agilent Technologies). Images were acquired at 20x magnification using a fluorescence microscope (Axioplan2, Zeiss).

### 5.6. Spatial transcriptomics on the Xenium platform

Formalin-fixed, paraffin-embedded tissue sections (5-µm-thick) were mounted onto Xenium slides. The tissue sections were subsequently deparaffinized and decrosslinked according to the Xenium Deparaffinization and Decrosslinking Protocol (Demonstrated Protocol CG000580, 10x Genomics). Xenium Prime pre-designed probes were applied to hybridize to the target RNA. Following hybridization, an RNase treatment was performed to release the RNA strands from the priming oligos. A polishing step was then carried out prior to the addition of the Xenium Prime 5K Human Pan-Tissue & Pathways probe set. These probes were hybridized to the target RNA, followed by ligation and enzymatic amplification, generating multiple copies of each RNA target, as described in the Probe Hybridization, Ligation, and Amplification User Guide (User Guide CG000760, 10x Genomics). Cell segmentation reagents were subsequently applied to label nuclei, cell membranes, and cytoplasmic compartments, providing the necessary inputs for automated, morphology-based cell segmentation. The prepared Xenium slides were then loaded onto the Xenium Analyzer for imaging and analysis, following the Decoding and Imaging User Guide (User Guide CG000584, 10x Genomics). Instrument software version 3.4.1.0 and analysis software version 3.3.0.1 were used for this workflow.

### 5.7. Immunofluorescence Staining on Xenium-processed slides

Cover slips of the Xenium-processed slides were removed in 1x PBS at 37°C. After a quick wash, HIER was performed using ready-made TRIS-EDTA, pH 9.0 (ab93684, Abcam), for 30 minutes at 80°C in a decloaking chamber (Biocare Medical). Following HIER, the slides were first treated with Background Punisher (BP974H, Biocare Medical) at RT for 10 minutes, followed by an additional hour of blocking at RT with 10% Goat Serum (B40913A, Thermo Fischer).

Tissue sections were incubated with a primary antibody for endothelial cells (CD31 (1:50), DIA-310, Dianova; Alexa Fluor 568 Conjugated IsolectinB4 (1:25), I21412, Thermo Fischer; and biotin-conjugated HABP (1:50), AMS_HKD_BC41 Amsbio) in 3% BSA in PBS overnight at 4°C. The next day, after a washing step, tissues were incubated with secondary antibody (Goat anti-Rat IgG (H&L) - Cyanine5 (1:200), and streptavidin-Cy7 (1:100), both from Thermo Fischer) in 3% BSA in PBS for an hour at RT. Nuclei were stained using DAPI (1:200, MBD0015, Sigma Aldrich). Images were taken at 40x magnification using a VS200 imaging scanner (Olympus).

### 5.8. Hematoxylin and Eosin (HE) Staining on Xenium-processed slides

Following FS, the cover slips from Xenium-processed slides were removed as described above. The quencher was removed by immersing slides in 10 mM sodium hydrosulfite (157953, Sigma-Aldrich) according to the 10X Genomics’ protocol (CG000613-B, 10X Genomics). Subsequently, a standard H&E staining was performed on the slides using Mayer’s hematoxylin (01820, HistoLab) and 0.2% Eosin Y solution (01650, HistoLab). Tissues were dehydrated in increasing grades of alcohol and xylene. Cover slips were sealed using Pertex (00840, HistoLab) and images were taken at 40x magnification using Slideview VS200 slide scanner (Olympus).

### 5.9. Image processing and analysis

All the image processing and analysis were carried out on a workstation equipped with an Intel® Xeon® W-2133 CPU (3.60GHz, 6 Cores), 256 GB RAM and an NVIDIA Quadro P5000 GPU, with Windows 10 as operating system.

The analysis was performed with a combination of different functions in the software programs FIJI and Amira (Thermo Fisher Scientific, Waltham, Massachusetts, US). It is important to note that, while FIJI is open source and freely available, Amira is a licensed software. While this software was used because of convenience and familiarity, functions similar to the ones employed in this work are also available in free software packages like FIJI itself, 3D Slicer, ilastik, ITK-SNAP, Dragonfly (free for academic users), in various more specific open source programming toolboxes for segmentation (e.g. scikit-image) or registration (e.g. elastix), as well as in multiple other licensed software tools.

#### 5.9.1. Morphology

A first evaluation was performed by scrolling through the volumes in FIJI or in Amira (using the “orthoslice” module). For a more precise interpretation, as well as for arbitrary selection of sectioning planes as shown in Figure 3, the “slice” module in Amira allowed easy rotation of the slicing direction.

The first semi-transparent volume rendering (Figure 2-A), prior to any segmentation, was done using the “volren” module in Amira.

All the segmentation was performed in Amira. First, the volumes were corrected for local tomography artefacts (which manifest as grayscale gradients throughout the volume) by estimating such artefacts with a second order polynomial fit (using the “background image” module) and dividing the original data with the estimated background. A median filter with a 3×3×3 kernel was then applied, to reduce image noise while still preserving edges.

Segmentation of the hearts was achieved using the “magic wand” tool, which performs a region growing operation in 3D from a manually placed seed point; the region growing was restricted to a certain mask, set to allow grayscale values pertaining to tissue and exclude the ones associated to the paraffin embedding. The segmentation was then cleaned with an iteration of morphological closing. In the cases where they had not been surgically removed, also parts of the lungs or the thymus were sometimes picked up by the “magic wand” tool if they were touching the heart; these were removed from the segmentation using the “lasso” tool, which allows to add or exclude voxels based on manual selection in 3D.

The segmentation pipeline up to this point would normally require around 30 minutes per heart, depending on the amount of extra tissue that had to be removed with the “lasso” tool, and on its positioning relative to the heart (a lot of tissue in very close proximity to the heart required a more careful selection).

In order to better visualize the different heart chambers in a 3D rendering, the pericardium was also excluded from the segmentation. This was achieved with a combination of morphological opening and then manual adjustments with “lasso” tool and “draw” tool. Similarly, to improve the appearance of the renderings, residual blood that had been included by the “magic wand” tool was removed from the ventricles and the atria. This was done with the “texture classification” tool, exploiting the fact that the residual blood had a grainier structure when compared to the rest of the segmented tissue. These cleaning steps, in particular the careful manual adjustments needed to completely remove the pericardium, added significant extra time (in the order of hours) to the segmentation.

The coronary circulation was segmented by first closing and filling the heart segmentation, and then using the “magic wand” tool restricted to the areas inside the heart and to an appropriate grayscale mask. Starting from the cleaned segmentation of the previous step, this further step was relatively quick to perform, although some manual adjustments were required in spots where the coronary vessels were occluded by residual blood.

#### 5.9.2. Phenotyping

The *Adamts6*-deficient heart was segmented with the same methods described in the previous section. In order to extract the luminal volumes of the cardiac chambers, the “compute ambient occlusion” module (based on ^51^) was applied in Amira to the heart segmentation. From the resulting ambient occlusion image, the cardiac chambers were extracted with the “magic wand” tool. In case of connections, they were then separated into distinct labels using the “lasso” tool. They were finally visualized either as semi-transparent volume renderings (all of Figure 3), or as semi-transparent mesh surface renderings (Figure 3-E and 3-F).

#### 5.9.3. Integration of histology and immunofluorescence

The pipeline that was used in this study is outlined in Figure 4 and can be considered as being formed of three main groups of operations.

The first group of operations has the purpose of preparing the data for the subsequent steps. The microscopy scans were first manually cropped so that they would roughly match the area that was scanned with PC-µCT. They were then converted from RGB to grayscale in FIJI, trying to obtain a relative contrast between the different types of tissue as similar to the one contained in the PC-µCT scans as possible. To this end, the RGB images were split using FIJI into their red, green and blue channels, or alternatively in their hue, saturation and brightness channels. The channel in which the contrast looked more like the PC-µCT scans was selected for grayscale conversion. This selection depended on the staining type, but once established a first time it was kept consistent for all sections of the same type. In the case of H&E staining, the green, saturation or brightness channels were all viable options (the red and blue channels instead would enhance the contrast for the cell nuclei, information that is not provided by PC-µCT at the employed resolution). Among those, the saturation channel was selected as it also reduced the artifacts caused by dust on the microscope’s lens. To save on computing time during the next step, the grayscale histological image was then binned to have the same pixel size as the PC-µCT data.

After data preparation, the second group of operations forms the first registration step, which was performed in Amira. Here, the grayscale binned histology and the PC-µCT data were loaded within the same 3D environment. The position of the histology relative to the PC-µCT was at first initialized manually, by dragging it approximately to the corresponding position in the PC-µCT volume (mostly caring that the data were not flipped and that rotation and height in the stack were reasonably close to the expected ones, but without dedicating a lot of time to fine adjustments). An automatic affine registration was then performed using the “register images” module. Since this module does not natively support 2D to 3D registration, the histology slice was first converted to a 3D format by simply duplicating it one time along the direction normal to the plane. The “register images” module was then applied, having the histology as moving data and the PC-µCT as static data. The module was run several times, each time with a progressively smaller initial step size but also larger number of degrees of freedom. In all the steps, the normalized mutual information metric and the “quasi-Newton” optimizer (with a final step size always left at the default value of 1/6 of the voxel size) were used. In the first run, the module was set to perform only a rigid registration, only on 2×2×2 binned data (i.e. ignoring the finest resolution), and with an initial step size of 20 for the optimizer (sometimes increased to even larger values after inspection of the results). The outcome was then visually assessed by comparing the histology image with the newly-found corresponding plane in the PC-µCT data. If no features were recognizable, the registration was reverted and the “register images” module was run again with a larger optimizer initial step size, or possibly the manual initialization was re-performed before running the module again. If similar features were recognizable, with differences mostly being due to different scaling, the “register images” module was run a second time (without undoing the results of the previous step), while enabling isotropic scaling as an additional degree of freedom beside rigid transformation, and reducing the optimizer initial step size (typically to a value of 4). Lastly, the module was re-run one final time (on top of the previous results), enabling anisotropic scaling beside isotropic and rigid transformation, using full-size resolution instead of binning, and reducing the initial step size even further (typically to a value of 1). After the previous steps were run for the binned histological image, the same affine transformation was also applied to the unbinned image (the scaling was automatically handled by the program after correctly setting the voxel sizes of binned and unbinned data).

At this point of the pipeline, the match between histology and PC-µCT typically looks acceptable at a low magnification, but is clearly far from a 1-to-1 match when evaluated at a zoomed-in level. This is due to local deformations of the tissue, which cannot be fixed with the global transformations performed so far. These deformations are addressed by the third group of operations in the pipeline, which consists of a 2D-to-2D elastic registration. From the previous step, the PC-µCT slice that matched the position of the histology was interpolated onto the higher-resolution grid of the unbinned histological data. This interpolated PC-µCT slice and the unbinned grayscale histology were then used as inputs for the bUnwarpJ plugin ^52^ in FIJI. The plugin performs 2D image registration based on elastic deformations represented by B-splines. The histology image was set as “source” (i.e. moving data) and the PC-µCT was set as “target” (i.e. static data), and the registration mode to “mono” (i.e. only from source to target, no need for bidirectional registration as we assumed the PC-µCT data as ground-truth). The plugin offers an option to perform a subsampling of the data before computing the registration; the transformation is then applied to the unbinned data. To reduce computing time, this option was used with a subsampling factor of 1, i.e. a 2×2 binning. The “initial deformation” and “final deformation” parameters, which regulate the coarsest and finest scale of the splice deformation field, were set to “fine” and “super fine” respectively (corresponding to 4×4 intervals and 16×16 intervals in the B-spline grid). The other parameters were left as defaults (“image weight” = 1.0, “consistency weight” = 10.0, “stop threshold” = 0.01, all the other weights were left at 0), and the option to save the transformation was enabled. After processing the grayscale histology, the saved transformation was then used to process the RGB original version. The output was loaded into Amira and positioned into the correct 3D orientation by applying to it the same affine registration obtained from the previous step. It was then possible to visualize it in the 3D environment together with renderings and virtual slices based on the PC-µCT data, as described in the previous section.

In the case of FS data, the only difference was in the pre-processing step, during which, in order to extract grayscale morphological information similar to PC-µCT, the DAPI staining image was isolated and heavily smoothed with a Gaussian filter of σ = 7 to fill out the space between the cell nuclei. The same steps described above were then performed and applied to the complete FS images.

#### 5.9.4. Integration of spatial transcriptomics

For the spatial transcriptomics data, the same registration pipeline as in the previous section was applied. The grayscale image used for morphological information was the “Interior – RNA” image (18S rRNA) that is acquired as part of the Xenium protocol for automatic cell segmentation.

After registration, the same transformations could be applied to any other image, such as the individual transcripts as shown in Figure 7 or clusters as shown in Figure 8. The fluorescent staining and H&E images acquired after ISS had to be registered as well, since the processing had deformed the tissue structure. The 2D-to-2D registration was performed using Amira and bUnwarpJ in a similar way as the 2D-to-3D registrations.

#### 5.9.5. Laboratory-based PC-µCT

The SRPC-µCT and LBPC-µCT datasets were registered to each other using the “register images” module in Amira selecting affine transformation with the mutual information metric. The same virtual slice was then extracted from all the volumes.

## Supporting information

Video_clip_1

Video_clip_2

## 6. Acknowledgements

We acknowledge the Paul Scherrer Institut, Villigen, Switzerland, for provision of synchrotron radiation beamtime at the TOMCAT beamline X02DA of the Swiss Light Source (proposal numbers 20201620 and 20191567).

We acknowledge SOLEIL for provision of synchrotron radiation facilities and we would like to thank Jonathan Perrin and Timm Weitkamp for assistance in using the ANATOMIX beamline under proposal 20240499.

This work was supported by the Swedish Heart-Lung Foundation (to KTL), the Swedish Research Council (to KTL), the Skåne County Council (to KTL), the Knut and Alice Wallenberg Foundation (to KTL) and a Paul G. Allen-American Heart Association Distinguished Scientist award (17DIA33820024 to SAA). NP was supported for part of his work by a scholarship from the Tegger Foundation. ACM is partially funded by AMBER Grant agreement ID: 101126665 (to KTL). The AMBER program is a Marie Skłodowska-Curie Actions Program within the European Commission MSCA framework. Funded by the European Union. Views and opinions expressed are, however, those of the author(s) only and do not necessarily reflect those of the European Union or MSCA. Neither the European Union nor the MSCA can be held responsible for them.

## 7. Author contributions

NP, KTL, KT conceived the study and designed the experiments. KTL, SAA and MB provided funding and resources. KIG and TJM performed animal experiments and prepared the tissue samples. NP, KTL, AB, TJF, ELagervall, ELampei, TD, RK and MB performed the synchrotron-based PC-µCT experiments. JR performed the laboratory-based PC-µCT experiment on the commercial scanner. RK and TD performed the laboratory-based PC-µCT experiment on the custom-made scanner. KIG, ELampei and KTL performed histological sectioning and staining for H&E and fluorescent staining. TJF and ELagervall performed the sectioning for in-situ sequencing. KT and RS performed the in-situ sequencing experiments. ACM performed fluorescent staining and H&E after in-situ sequencing. NP performed the data analysis. NP and KTL wrote the manuscript. All authors contributed to the final version of the manuscript.

## Notes

### Competing Interest Statement

The authors have declared no competing interest.

