## Supplemental material for "Efficient murine cardiac phenotyping by combining synchrotron-based phase-contrast micro-CT, histology, immunofluorescence and spatial transcriptomics"

**Figure S1** – comparison of actual field-of-view (FoV) between SRPC-µCT and LBPC-µCT. **A** – full FoV of SRPC-µCT, as shown in Figure 9-A. **B** – LBPC-µCT cropped to the FoV of SRPC-µCT, as shown in Figure 9-B. **C** – actual FOV of the LBPC-µCT with the commercial scanner. The area corresponding to A and B is highlighted with a white dashed outline.

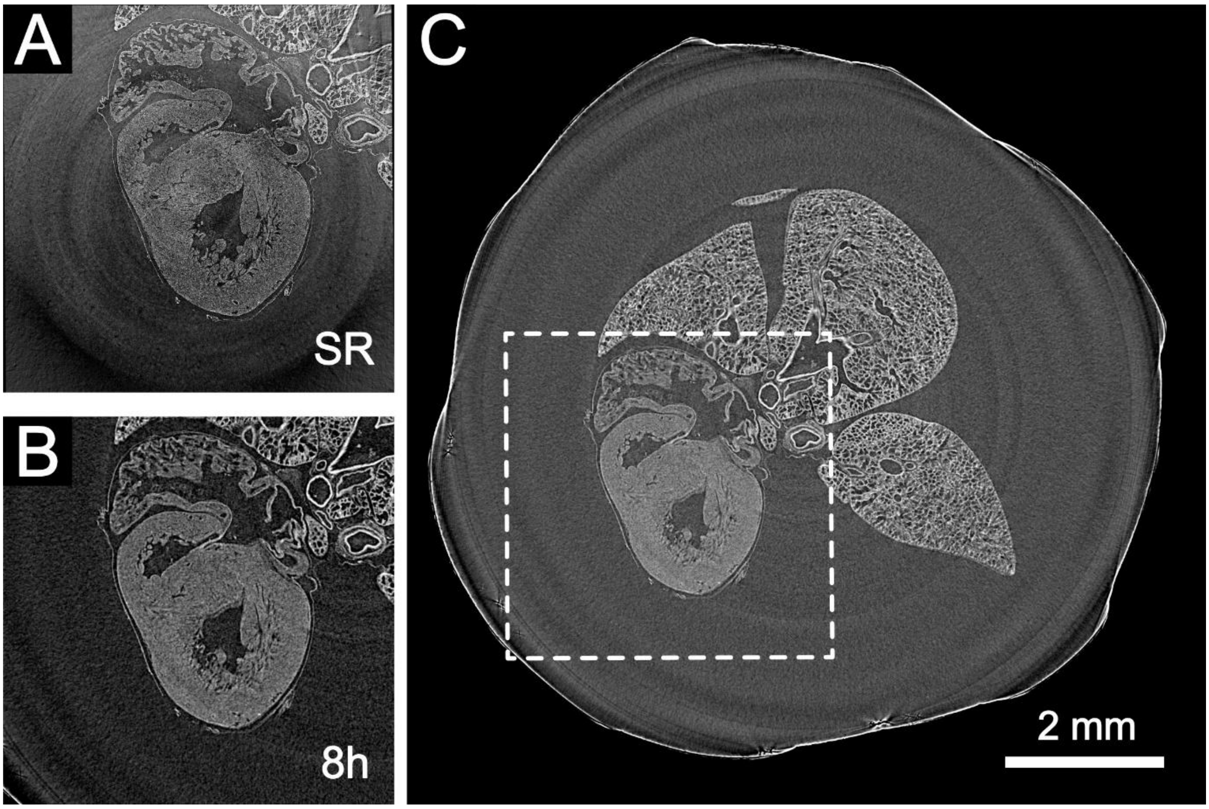

**Figure S2** – comparison between commercial and custom-made LBPC-μCT. **A** – full FoV of the custom-made LBPC-μCT (the solid white outline shows the full extent of it, as local tomography artefacts cause part of the image to be completely dark if the contrast is optimized for the tissue). The full image would be the FoV of the commercial system with the settings used in this work, as shown in Figure S1-C. **B** – Cropped commercial LBPC-μCT scan, with inset showing smaller details. Reproduced from Figure 9-B for convenience. **C** – LBPC-μCT scan from the custom-made setup cropped to the same region shown in Figure S2-B and Figure 9. Despite using a cheaper and less powerful x-ray source, the image quality is comparable in even less time if one looks at the larger features. This is probably due to the higher efficiency of the (more expensive) photon counting detector employed in the custom-made system. When looking at finer details however, the smaller coronary arteries are noticeable but less pronounced (orange arrow), almost in-between the 8h and 2h scans of the commercial system. The larger coronaries are well visible in both scans (green arrow).

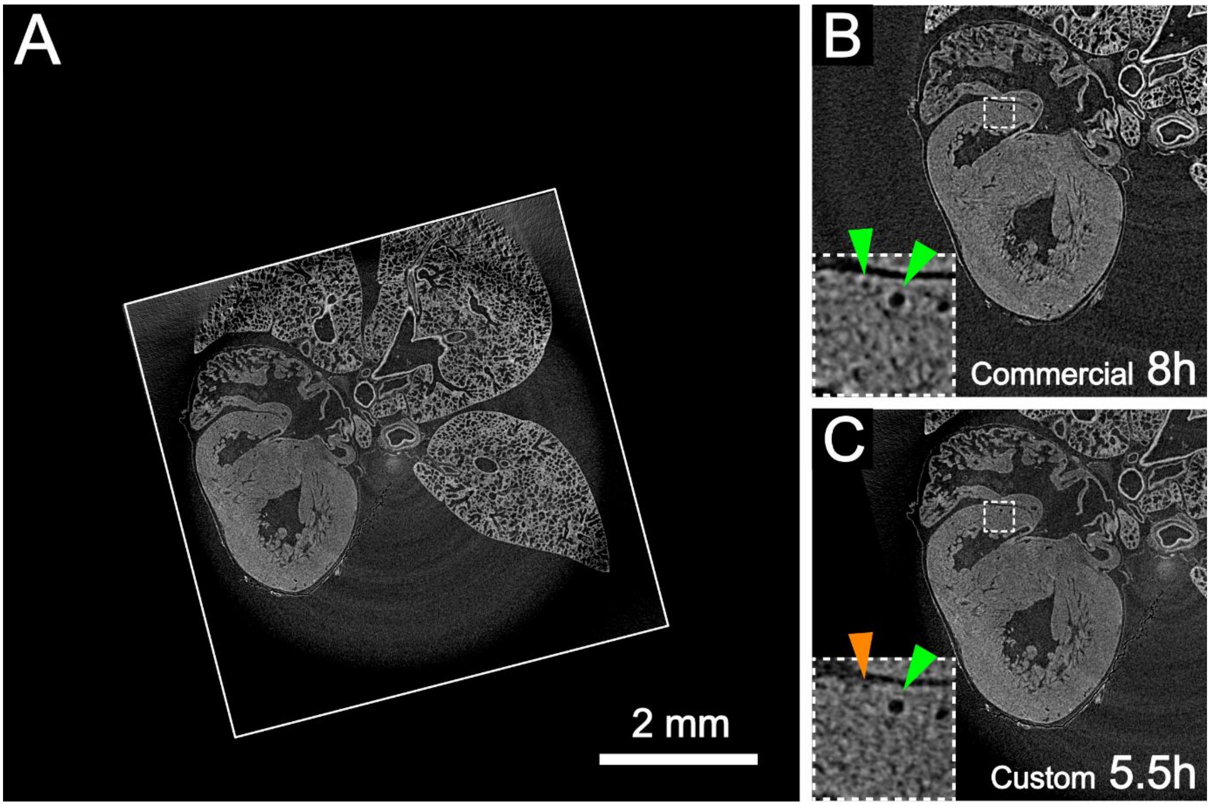
